# Selective kinase inhibition shows that Bur1 (Cdk9) phosphorylates the Rpb1 linker in vivo

**DOI:** 10.1101/507251

**Authors:** Yujin Chun, Hyunsuk Suh, Yoo Jin Joo, Gaelle Batot, Christopher P. Hill, Tim Formosa, Stephen Buratowski

## Abstract

Cyclin-dependent kinases play multiple roles in RNA polymerase II transcription. Cdk7/Kin28, Cdk9/Bur1, and Cdk12/Ctk1 phosphorylate the polymerase and other factors to drive the dynamic exchange of initiation and elongation complex components over the transcription cycle. We engineered strains of the yeast Saccharomyces cerevisiae for rapid, specific inactivation of individual kinases by addition of a covalent inhibitor. While effective, the sensitized kinases can display some idiosyncrasies, and inhibition can be surprisingly transient. As expected, inhibition of Cdk7/Kin28 blocked phosphorylation of the Rpb1 C-terminal domain heptad repeats at Serines 5 and 7, which are known target sites, but also at Serine 2. Consistent with our previous results using gene deletions, Cdk12/Ctk1 is the predominant kinase responsible for Serine 2 phosphorylation. Phosphorylation of the Rpb1 linker enhances binding of the Spt6 tSH2 domain, and here we show that Bur1/Cdk9 is the kinase responsible for this modification in vivo.

## Introduction

Phosphorylation provides a powerful mechanism for rapidly and reversibly affecting protein functions and interactions, and a very large percentage of eukaryotic proteins are phosphorylated in vivo. Accordingly, there is great interest in chemical kinase inhibitors, both for scientific studies and as potential drugs. The greatest challenge is to create compounds with sufficient specificity to overcome the structural similarities among kinase active sites. One powerful strategy is the “bump-hole” system, in which a “gatekeeper” residue in the target kinase ATP-binding pocket is mutated to create space for an ATP mimic carrying a bulky side group that blocks its binding to other kinases (1). Such mutated kinases are referred to as “altered specificity” (AS). This approach can be extended by engineering a reactive group on the inhibitor such that it covalently links to a specifically-positioned cysteine found on the target kinase. Such covalent inhibitors are irreversible, which can produce more complete kinase inactivation. Using these approaches, an “irreversibly sensitized” (IS) kinase allele can be created with two point mutations (2–4).

A great deal of effort has gone into understanding the functions of the cyclin-dependent kinases (CDKs) involved in transcription by RNA polymerase II (reviewed in (5–7)). Cdk7/Kin28 assembles into the pre-initiation complex (PIC) as part of the basal transcription factor TFIIH. It phosphorylates the C-terminal domain (CTD) of polymerase subunit Rpb1, primarily on Serine 5 of the repeated consensus sequence YSPTSPS. This phosphorylation triggers several events near 5′ ends of genes: it dissociates the Mediator complex that chaperones RNApII into the PIC, while creating binding sites for the mRNA capping enzyme, the Nrd1/Nab3 early termination complex, and the Set1/COMPASS histone methyltransferase. As RNApII proceeds into elongation, two additional CDKs come into play. Cdk9/Bur1 phosphorylates the C-terminal region (CTR) of the Spt5/DSIF elongation factor, which in turn promotes binding of the PAF1 complex (PAFc), another important elongation factor. In some eukaryotes, Cdk9/Bur1 also phosphorylates the negative elongation factor NELF to reverse its inhibitory effects. Cdk9/Bur1 may also phosphorylate Serine 2 of the Rpb1 CTD at a low level during early elongation (8), although other experiments suggest it also targets Ser5 (reviewed in (6)). In any case, the vast majority of CTD Ser2 phosphorylation is due to Cdk12/Ctk1 during productive elongation. Ser2 phosphorylation promotes binding of the histone methyltransferase Set2 and multiple polyadenylation/termination factors to RNApII complexes as they proceed downstream (5–7).

AS alleles have frequently been used to inhibit these transcription-related kinases in budding and fission yeast (8–17) as well as mammalian cells (18–20). Chemical inhibitors are available that target the native mammalian kinases with varying levels of specificity (21–23). Importantly, the Ansari lab (4) demonstrated that a Kin28-IS inhibition was much more effective than Kin28-AS inhibition in yeast cells, possibly because powerful drug efflux channels limit the concentrations of inhibitors that can be maintained in vivo. Therefore, AS allele inhibition experiments have produced conflicting conclusions and failed to reveal effects seen using IS allele inhibition. Here we extend the IS approach to Bur1/Cdk9 and Ctk1/Cdk12.

We created double point mutations that confer sensitivity to the chemical CMK, which has both a “bump” that can be accommodated by a “hole” mutation in the ATP binding pocket, and a functional group that covalently crosslinks to an engineered cysteine near the active site (2, 3). We find that the double mutants are expressed and functional, although they may not have full wild-type activity or stability. Inhibition in vivo was very rapid but surprisingly transient in liquid cultures, demonstrating the need for choosing an appropriate time point after CMK addition. As predicted, Kin28/Cdk7 inhibition reduced Ser5P and Ser7P, while Ctk1/Cdk12 inhibition blocked Ser2P. In contrast to most previous reports (see Discussion), we found that Ser2P was also strongly blocked upon Kin28 inhibition, indicating clear sequential dependence of the two marks. Bur1 inhibition had less effect on CTD phosphorylation, supporting our earlier findings that Bur1/Cdk9 is not a major Ser2P kinase (24). However, we discovered that Bur1/Cdk9 phosphorylates the Rpb1 linker region, a domain that lies between the RNApII body and the CTD. Phosphorylation of specific Rpb1 linker residues enhances binding of the Spt6 tandem SH2 (tSH2) domain (25, 26), indicating that Bur1/Cdk9 activity is important for linking both elongation factors Spt5/DSIF and Spt6 to the elongating RNApII.

## Results

### Creation of irreversibly sensitized kinase alleles

Cohen et al. created the covalent kinase inhibitor CMK as an inhibitor of RSK and PLK family kinases. This molecule is an adenine-like pyrrolopyrimidine derivative that carries both a bulky “bump” constituent and a chloromethylketone (hence the name CMK) group that covalently links to a reactive cysteine found in this family of kinases (2, 3). Rodriguez-Molina et al. (4) showed that CMK sensitivity could be conferred on a Kin28 mutant combining “hole” (L83G) and cysteine (V21C) mutations.

Using alignments of the Kin28/Cdk7, Bur1/Cdk9, and Ctk1/Cdk12 sequences, we designed corresponding “hole” and reactive cysteine mutants for Bur1 and Ctk1 (**Fig 1A**). The previously described Bur1 AS mutation L149G (8, 27) creates the “hole”, while changing valine 74 to cysteine (V74C) creates the covalent linkage site. Similarly, the combination of F260G and V197C mutations are predicted to create a Ctk1-IS protein. The mutated genes were introduced into yeast using plasmid shuffling, and protein expression levels were tested using a triple hemagglutinin (HA3) tag introduced onto the C-terminus. Immunoblotting showed that Kin28-IS and Ctk1-IS proteins were expressed at levels similar to their wild-type counterparts (**Fig 1B**). It should be noted that, unlike *Kin28* and *BUR1, CTK1* is a non-essential gene, so retention of the Ctk1-IS plasmid requires use of selective growth medium. Bur1 levels are significantly lower than the other two kinases, necessitating a longer exposure for detection (**Fig 1B**, bottom panel). Bur1-IS protein expression was lower than wild-type, indicating that the dual mutations affected its stability. Each single mutation also caused some reduction, apparently contributing additively in the double mutant (**Sup Fig 1A**). Bur1-IS levels could be boosted above normal wild-type levels by expressing the mutant on a high-copy plasmid (**Sup Fig 1B**), but these cells grew noticeably slower than wild-type. Treatment of cells with CMK did not affect levels of any of the kinases (**Fig 1B**).

**Figure 1.**
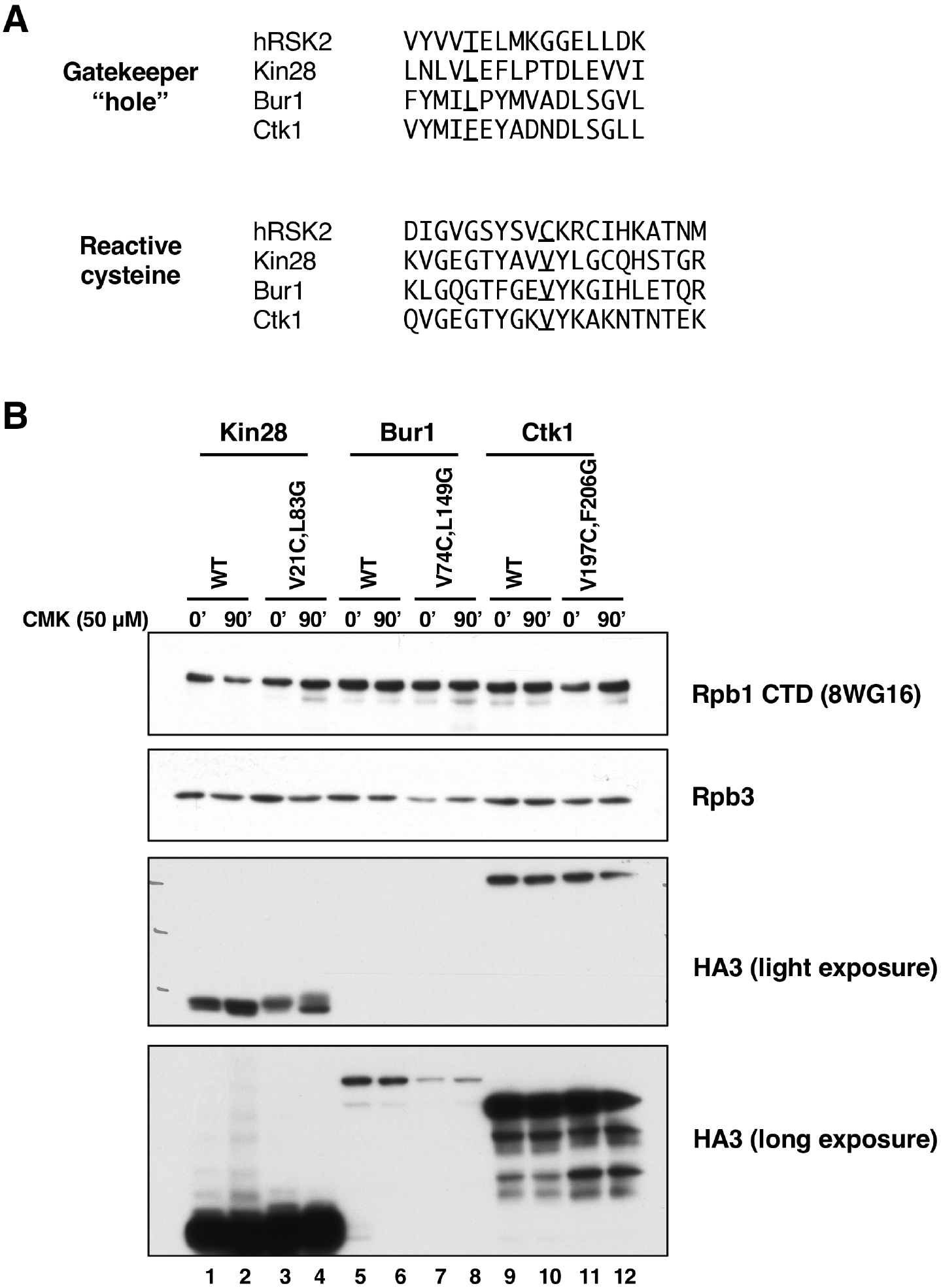
Construction of irreversibly sensitized (IS) kinase strains. A. Sequence alignments of IS mutant positions (2–4) in human RSK2, Kin28, Bur1, and Ctk1 kinases. B. Protein expression levels of wild-type and IS mutant kinases, before (0) or after (90) treatment with 50 μM CMK. Anti-HA blot shows epitope-tagged kinases at the expected sizes of Kin28 (35 kDa), Bur1 (74 kDa), and Ctk1 (61kDa). Rpb1 and Rpb3 are two RNA polymerase II subunits control bands. Strains used: YSB3216 (Kin28 WT), YSB3221 (Kin28 V21C, L83G), YSB3229 (Bur1 WT), YSB3232 (Bur1 V74C, L149G), YSB3235 (Ctk1 WT), YSB3237 (Ctk1 V197C, F206G).

Growth rates of IS strains were similar to a wild-type at 30°C, indicating that the mutated kinases were functional. Bur1-IS grew slightly slower than WT at 16°C, possibly reflecting the reduced protein levels. While wild-type cells were unaffected, both Kin28-IS and Bur1-IS cells showed strong inhibition of growth by CMK in plate spotting assays (**Fig 2**). Ctk1-IS cells produced smaller colonies on CMK plates, particularly at 16°C, consistent with the phenotypes of ctk1Δ cells. These results indicate that CMK enters cells and inhibits the target kinases.

**Figure 2.**
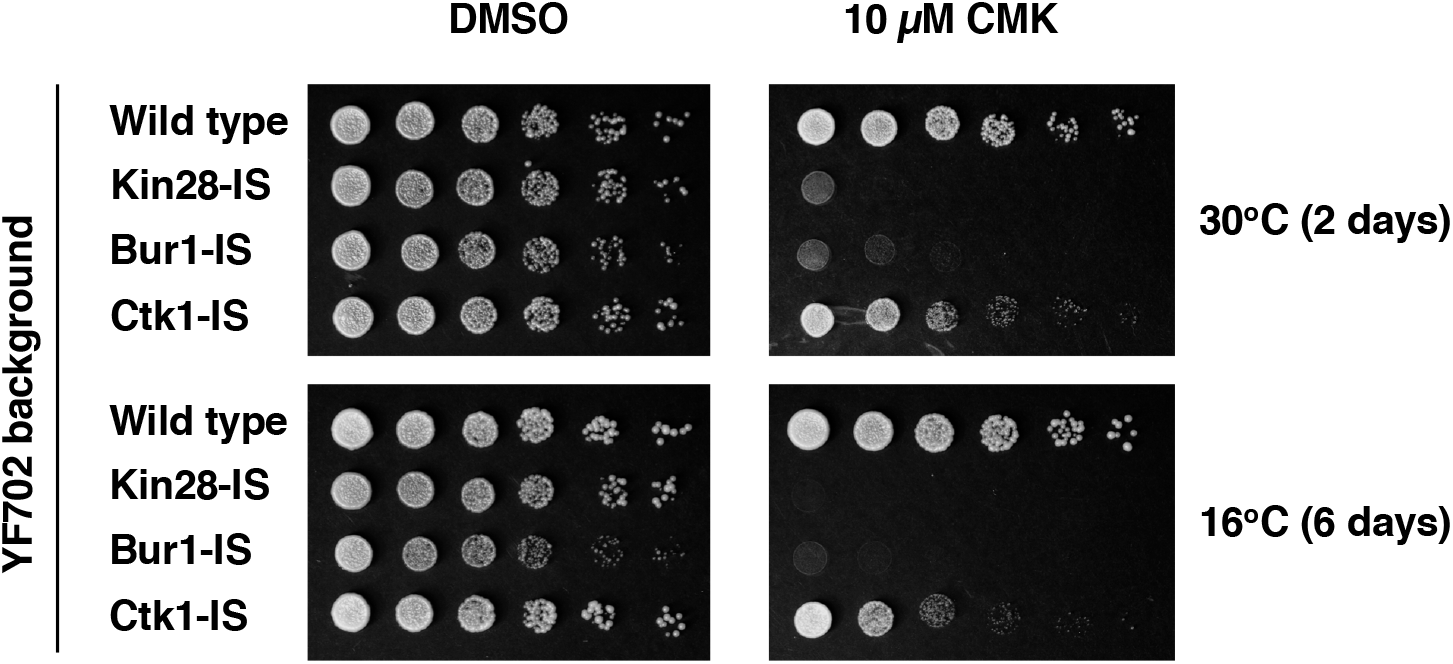
Growth inhibition of IS strains on CMK plates. Strains YF702 (Wild Type), YSB3356 (Kin28-IS), YSB3419 (Bur1-IS), YSB3444 (Ctk1-IS) were spotted on YPD plates containing 10 μM CMK. Each row shows three-fold dilutions. Plates were photographed after the indicated number of days at the indicated temperature.

### Inhibition by CMK is effective but transient in liquid cultures

Given the clear growth inhibition on CMK plates, we were surprised to find that growth of the IS strains in liquid media was not strongly affected by CMK. This observation contrasted with a previous study using Kin28-IS in a different strain background (4). In our hands, concentrations between 1 and 50 μM CMK failed to completely arrest growth in rich or minimal media liquid cultures. To probe this discrepancy and to validate that CMK was effectively inhibiting the kinases, CTD phosphorylation was monitored using immunoblotting (**Fig 3A**).

**Figure 3.**
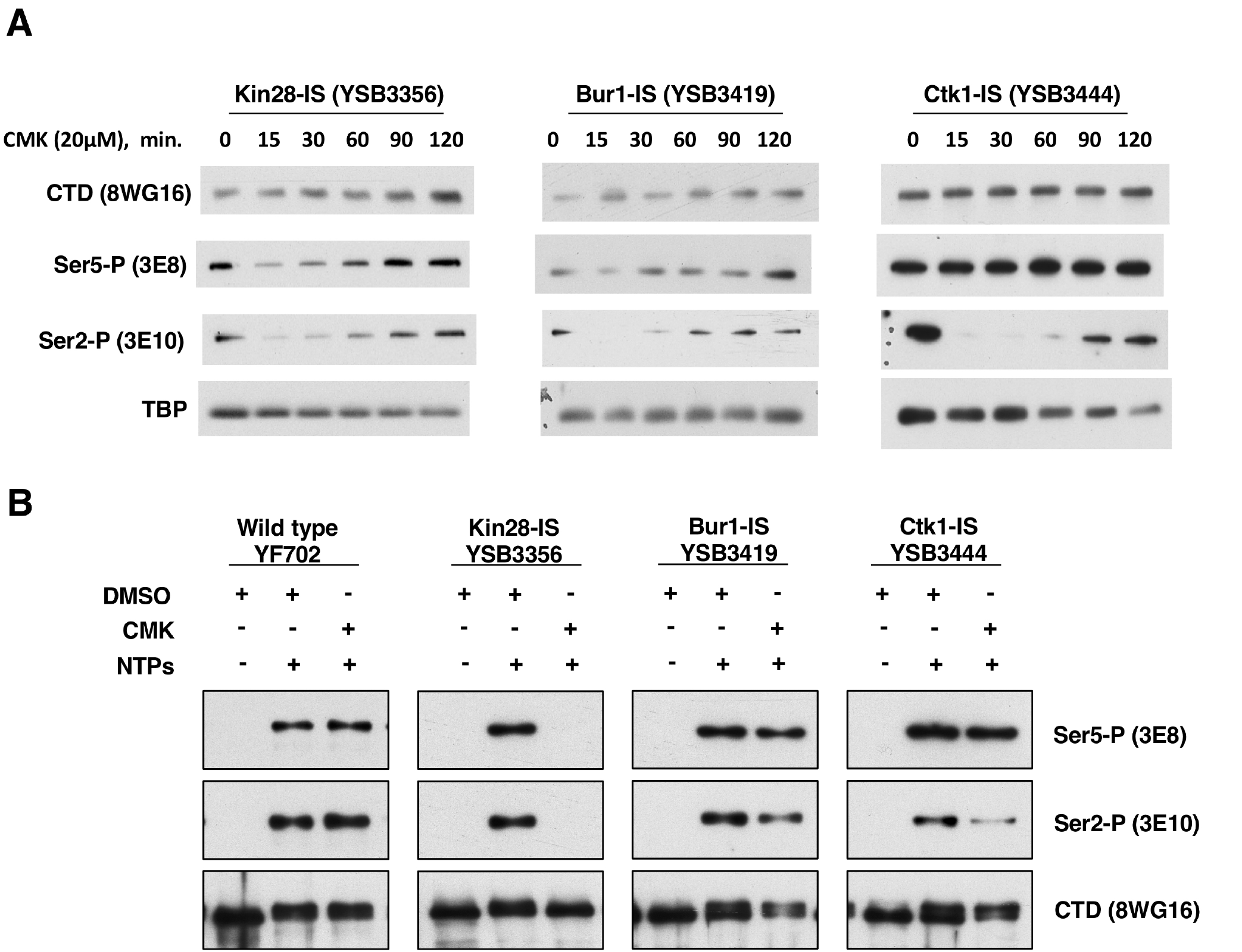
Inhibition of CTD phosphorylations in the IS kinase strains. **A.** Time course of CTD phosphorylation levels during CMK inhibition in vivo. The indicated yeast strains were grown to mid-log phase (OD 1.0) at 30°C. Immediately after taking a sample to serve as the zero time point, CMK was added to 20 μM final concentration. At each indicated time point, samples were taken and processed for immunoblotting as described in the Methods section. Blots were probed with the indicated antibodies. **B.** Transcription complexes assembled in vitro. Yeast nuclear extracts were prepared from the indicated strains and incubated with DNA template carrying five Gal4 binding sites upstream of the CYC1 core promoter and a G-less cassette. Immobilized templates were isolated and analyzed by immunoblotting. The first lane in each set shows RNApII complexes formed in the absence of NTPs. The second and third lanes show complexes formed upon addition of ATP, UTP, and CTP for four minutes, with the third lane showing the effect of CMK inhibition.

### Inhibition of each kinase gave a different response

Notably, time courses revealed that CMK inhibition of all three kinases was very rapid, but transient. For example, a 15-minute CMK treatment of Ctk1-IS cells caused very strong loss of Ser2P but no effect on Ser5P, as expected for the major Ser2 kinase. However, Ser2P showed partial recovery by 90 minutes (**Fig 3A**). As assayed by multiple CTD antibodies, Ctk1-IS inhibition for 15 minutes mimics phosphorylation patterns seen in a *ctk1Δ* strain (**Fig 4**). Inhibition of Bur1 for 15 minutes also reduced Ser2P, although not as completely as Ctk1 inhibition (**Fig 3A, 4**). Ser2P recovery in the Bur1-IS strain was faster and to a greater extent than Kin28 (**Fig 3A, 4**). These results show that the time frame for maximal effect of these covalent inhibitors must be established and interpretations should be limited to that window. Given that the S. cerevisiae cell cycle in rich medium is about 90 minutes, this transient response explains why CMK did not arrest the growth of liquid cultures.

**Figure 4.**
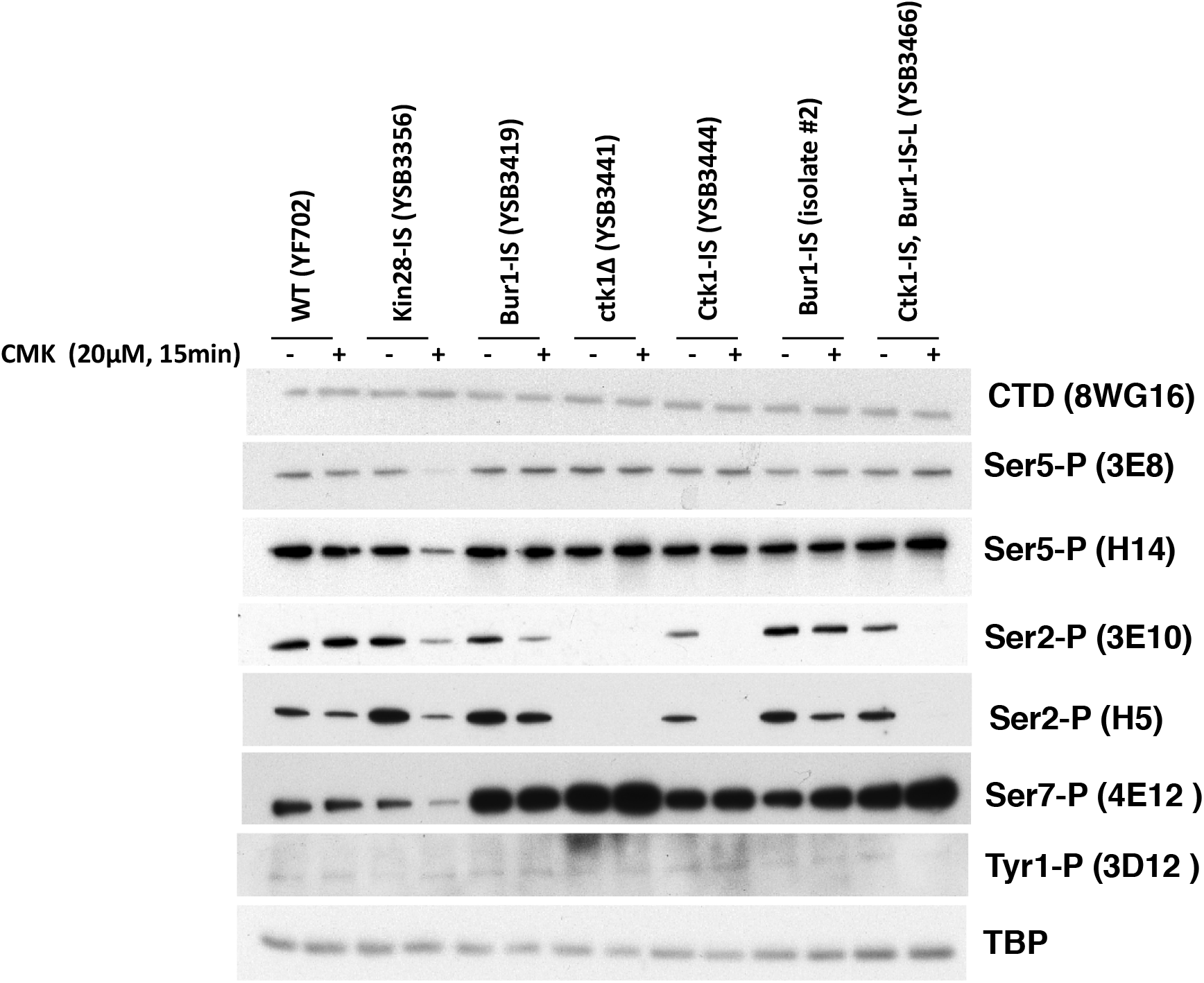
Analysis of IS kinase inhibition with different CTD antibodies. The indicated yeast strains were analyzed before or after 15 minutes of inhibition with 20 μM CMK. Blots were probed with the indicated antibodies. TATA-Binding Protein (TBP) is shown as a loading control. Note that all blots were developed with Pierce Supersignal West Pico Chemiluminescent Substrate, except for 4E12 (Ser7-P) and 3D12 (Tyr1-P), which required the Femto Maximum Sensitivity Substrate. No signal above background was detected with 6G7 (Thr4-P, not shown).

As expected, treatment of the Kin28-IS strain with CMK led to a significant, albeit incomplete, drop in CTD Ser5P. This was seen with both 3E8 and H14 antibodies (**Figs 3A and 4**). Ser7P, another known target site for Kin28/Cdk7, was similarly reduced (**Fig 4**). The remaining low level of these modifications may reflect incomplete Kin28 inhibition, Ser5P phosphorylation by a different kinase, or persistence of some Ser5P phosphorylated before CMK addition. As seen for Ctk1-IS and Bur1-IS, Kin28-IS inhibition was also transient, with recovery apparent by 60 minutes after CMK addition. In contrast to Rodriguez-Molina et al (4) and multiple Kin28-AS studies (11, 12, 14, 15, 17), we found that Kin28-IS inhibition also strongly inhibited Ser2P formation, as assayed by either 3E10 or H5 antibodies (**Figs 3A and 4**).

We confirmed the effects of kinase inhibition using yeast nuclear extracts to assemble RNApII complexes on DNA templates containing five Gal4 binding sites upstream of the *CYC1* core promoter and a G-less cassette (28). The bead-immobilized templates were incubated with IS strain extracts treated with CMK or solvent DMSO. Complexes were recovered magnetically, and bound proteins were isolated and analyzed by gel electrophoresis and immunoblotting (**Fig 3B**). Pre-initiation complexes formed in the absence of NTPs have no CTD phosphorylation (first lane in each kinase set). Elongation complexes stalled at the end of the G-less cassette were formed by adding ATP, UTP, and CTP for four minutes (second and third lanes). As we previously reported (28), and in agreement with the in vivo results (**Fig 3A**), Kin28 inhibition in vitro blocked both Ser5 and Ser2 phosphorylation. In contrast, Ctk1 inhibition specifically reduced Ser2P, as did Bur1 inhibition to a lesser extent. As seen in vivo, neither Ctk1 nor Bur1 reduced Ser5P levels.

### Bur1 phosphorylates residues in the Rpb1 linker region

In addition to the Rpb1 heptamer repeats, phosphorylations have been detected on several serines and threonines in the Rpb1 linker region just N-terminal to the CTD (25, 26, 29). These residues are also likely substrates for cyclin-dependent kinases, as they are followed by proline. Like the CTD, the linker region is apparently flexible, as it was not apparent in earlier RNA polymerase II crystal structures. However, Rpb1 linker phosphorylation promotes binding of the Spt6 elongation factor, via contacts between the Spt6 tandem SH2 (tSH2) domain and phosphorylated Rpb1 residues T1471 and S1493 (25). Interestingly, the phosphate on T1471 combines with non-phosphorylated Y1473 to occupy the pocket that recognizes phospho-tyrosine in other SH2 proteins (25). A recent cryo-EM reconstruction of the mammalian elongation complex was able to dock a linker-tSH2 crystal structure into density near the Rpb6-Rpb7 interface (26).

The kinase responsible for these Rpb1 linker phosphorylations in vivo has not yet been identified. To address this question, an MBP-Rpb1 linker region fusion protein was phosphorylated with either Bur1 or Ctk1 and tested for in vitro binding to the Spt6 tSH2 domains using “far-western” blotting (**Fig 5A, Sup Fig 2A, B**). No binding was observed without phosphorylation of the linker (None and Mock lanes). Although Bur1 gave a stronger signal, both kinases promoted tSH2 binding, which was abolished when the phosphorylated residues were mutated (**Fig 5A**). Therefore, the Rpb1 linker sites can be phosphorylated by the Cdk kinases in vitro. In agreement, a recent paper showed that mammalian Cdk9 can phosphorylate the corresponding linker sites in vitro, although other kinases were not tested (26).

**Figure 5.**
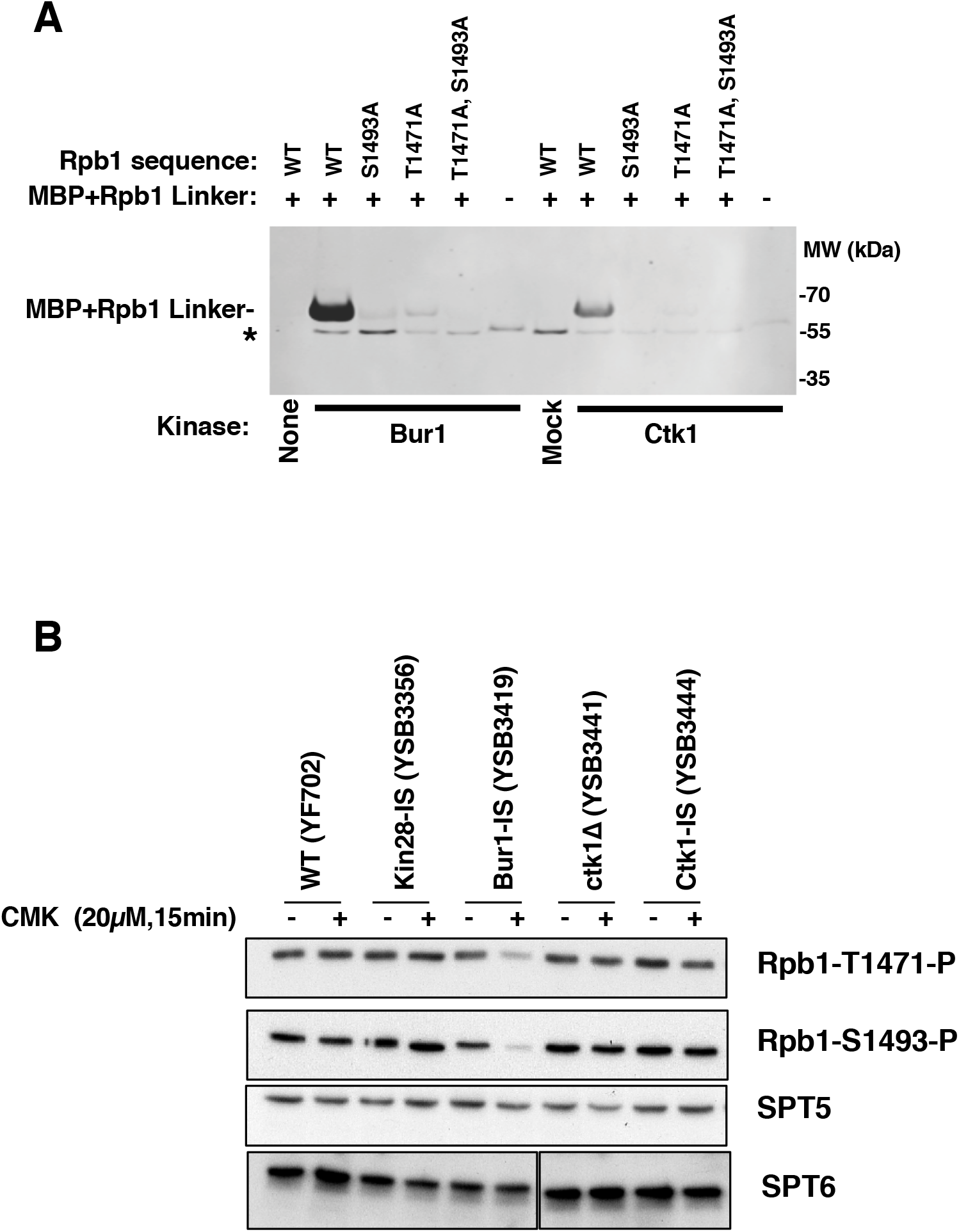
Bur1 phosphorylates Rpb1 linker region residues that mediate interaction with Spt6. **A.** Phosphorylation by Bur1 or Ctk1 enhances in vitro binding of Spt6-tSH2 to an Rpb1 linker peptide. MBP fusions to the Rpb1 linker (residues 1456-1511) were purified and incubated with His_12_-Bur1 or His_12_-Ctk1 partially purified from yeast, or the corresponding fractions from a mock purification from a strain without the affinity tags. Proteins were separated by SDS-PAGE, transferred to nitrocellulose, then probed with GST-Spt6 tSH2 and detected with antibodies against GST. Strong binding to the WT linker peptide was observed after treatment with Bur1, but this was diminished in mutants lacking the sites previously shown to be important for this interaction ((25); S1493 and T1471). Ctk1 also supported binding at a lower level. The asterisk denotes a species found in Bur1, Ctk1, or mock purifications that bound GST-Spt6 tSH2. **B.** The indicated yeast strains were analyzed before or after 15 minutes of inhibition with 20 μM CMK. Blots were probed with the antibodies specific for Rpb1 linker sites (see **Sup Fig 2**), or Spt5 and Spt6 as loading controls. Note that the Spt6 bands are all from the same exposure of a single blot, but intervening lanes were removed.

To identify which kinase phosphorylates the linker in vivo, extracts from the different IS kinase strains were immunoblotted with antibodies specific for the two phosphorylated residues that support Spt6-tSH2 binding (**Sup Fig 2C**). The results showed that Bur1/Cdk9 was the only required kinase. Both T1471 and S1493 phosphorylation were unaffected by Kin28-IS or Ctk1-IS inhibition, but were strongly reduced by CMK treatment of the Bur1-IS strain (**Fig 5B**). In contrast to CTD Ser2P, the linker phosphorylations were not reduced by Kin28 or Ctk1 inhibition, suggesting that neither kinase is required for Bur1/Cdk9 recruitment or activity on the linker.

## Discussion

To understand the role of an individual kinase, one must observe the effects of inactivating its function in vivo. There are multiple methods to accomplish this, each with its own advantages and disadvantages. Genes for non-essential kinases can be deleted, although care must then be taken that phenotypes don’t become masked by suppressing mutations accumulated during long term passaging. Temperature sensitive (TS) alleles can be very informative, but products of these genes are often partially defective at permissive temperatures or remain partially functional at non-permissive temperatures. Degron fusions can support normal function under permissive conditions, but loss of function may be slow or incomplete. Interpreting outcomes from these methods can also be complicated, as loss of the kinase protein can have secondary effects distinct from the loss of the kinase activity, such as destabilization of interacting factors. Chemical inhibitors can overcome these issues, as the protein remains present while kinase activity is ablated. However, this approach requires the availability of an inhibitor that completely inactivates the target kinase without affecting other kinases.

Here we applied the IS allele approach (2–4) to the three major transcription kinases, Kin28/Cdk7, Bur1/Cdk9, and Ctk1/Cdk12. This inhibitor approach significantly improves kinase specificity by combining the gatekeeper hole mutation with an additional cysteine substitution that allows irreversible inhibition through covalent linkage (**Fig 1A**). We find that CMK inhibition of IS alleles is effective and informative. However, the two sensitizing mutations can affect kinase function or protein levels in the absence of the inhibitor (**Fig 1B**). This is not unexpected, given that the “hole” mutation is in the ATP-binding pocket. We find that Ctk1-IS is expressed as well as wild-type Ctk1 protein, but lower levels of CTD Ser2P suggest activity is reduced in the mutant. For Bur1, the IS protein is expressed at reduced levels, although these can be boosted by increasing gene dosage. Despite the decrease in Bur1-IS protein, phosphorylation of targets in the Rpb1 linker region and CTD appear normal in the absence of CMK inhibitor, indicating an adequate level of Bur1 activity. The IS kinase strategy therefore provides an important probe for kinase function *in vivo*, but effects on protein expression and activity prior to addition of inhibitor must be monitored.

Although covalent modification of the kinases is presumably non-reversible, we found that the effects of CMK inhibition can be transient in yeast liquid cultures. Although growth of Kin28-IS or Bur1-IS on plates was completely arrested on solid media containing CMK (**Fig 2**), CMK generally did not arrest growth of liquid cultures. Immunoblotting showed that CTD phosphorylation was effectively inhibited immediately after CMK addition, but recovered by 60-90 minutes (**Fig 3**). Given that CMK covalently links to the target kinase, it seems unlikely that the inhibitor dissociates once bound. The very rapid recovery also rules out the outgrowth of rare inhibitor-resistant mutants. Most likely, CMK is depleted or inactivated as cells continue to synthesize more of the target kinase. Inhibitor binding is in kinetic competition with the efficient drug export pumps of yeast cells and the lability of CMK. Continued production of the IS kinase, perhaps by translation of existing mRNA, may eventually exceed the effective concentration of inhibitor. This effect likely explains the difference between liquid and solid media assays, since the latter starts with much lower density of cells and maintains a large reservoir of media where no cells are present. If so, repeated dosing with CMK might block recovery in liquid culture.

The previous characterization of Kin28-IS by Rodriguez-Molina (4) nicely demonstrated the utility of kinase inhibition, but we note several differences with our results. First, they found that CMK inhibited Ser5P as assayed by the IgM monoclonal antibody H14, but not using the IgG 3E8 monoclonal antibody. In our hands, CMK inhibits reactivity with both anti-Ser5P antibodies similarly (**Fig 4**). We also observed stronger Ser7P inhibition. Some of these differences may represent strain background differences. However, their experiments typically assayed cells after 60 minutes of CMK treatment. In our hands, significant recovery of CTD phosphorylation was apparent by 60 minutes, at which point our results more closely resemble theirs. In our strains, fifteen minutes was an optimal time for observing strong inhibition before any apparent recovery.

Another important observation is that in vivo inhibition of Kin28-IS produces strong inhibition of Ser2P, as well as Ser5P and Ser7P. The Ser2P drop was unexpected, as this phosphorylation was reported to be unaffected in multiple earlier studies using AS alleles of Kin28/Cdk7 in budding or fission yeasts (11, 12, 14, 15, 17), as well as in a paper using the same Kin28-IS allele used by us (4). Mammalian Cdk7-AS cells showed a partial reduction in both Ser5P and Ser2P (19, 20), while inhibition of mammalian Cdk7 with the covalent kinase inhibitor THZ1 strongly blocks Ser5P and Ser2P (21, 30). However, THZ1 also inhibits the Ser2 kinase Cdk12, and a recent paper using more specific Cdk7 and Cdk12 inhibitors again failed to see a Ser2P drop upon Cdk7 inhibition (31). In the in vivo systems, complete kinase inhibition is likely difficult to achieve and maintain (**Fig 3A**), but the dependence of Ser2P on Kin28/Cdk7 activity in yeast appears absolute in vitro using transcription on immobilized templates (28; see also **Fig 3B**).

Given the well-characterized in vitro specificity of Kin28/Cdk7 for CTD Ser5 and Ser7, our CMK results suggest that efficient Ser2 phosphorylation depends on Kin28 first phosphorylating either the CTD or some other substrate. There are several possible mechanisms that could explain this obligatory sequence of phosphorylations. One is that phosphorylation by Kin28 simply makes the CTD accessible to a Ser2 kinase. For example, CTD Ser5 phosphorylation releases Mediator (32), which may otherwise block Ser2 kinases. A second model is that Ser5P promotes binding or activity of Ser2 kinases. In vitro experiments suggest that yeast Ctk1 (33), metazoan Cdk12 (34), and mammalian Cdk9 (6) more efficiently phosphorylate Ser2 on a CTD “primed” by prior phosphorylation at Ser5 or Ser7. Supporting a recruitment model, Qiu et al (8) found that crosslinking of Bur1 to the *ARG1* gene was partially reduced when a *kin28-ts16* strain was shifted to non-permissive temperature. Although they did not actually show that Ser2P was reduced under these conditions, they speculated that Ser5P recruitment of Bur1 promotes Ser2P. Arguing against this model is our observation that Kin28-IS inhibition does not affect Bur1’s function in phosphorylating the Rpb1 linker region (**Fig 5B**). Furthermore, we find that Bur1 is still recruited to in vitro elongation complexes after Kin28-IS inhibition (28). Yet another possible linkage mechanism is that Kin28 acts as a CDK-activating kinase (CAK) to phosphorylate Ctk1 or Bur1. Although the Cak1 kinase carries out this function in budding yeast, Cdk7 phosphorylation of the Cdk9 T-loop has been reported in mammalian cells (19). Although the detailed mechanisms remain to be worked out, our demonstration that Kin28 activity is required for subsequent Ser2 phosphorylation helps explain the sequential nature of these two modifications.

Although in vivo inhibition of either Bur1 or Ctk1 caused a drop in CTD Ser2P, the loss in Ctk1-IS was far more complete (**Fig 4**). Liu et al. (27) and Qiu et al (8) observed little change in Ser2P upon inhibition of Bur1-AS unless it was combined with *ctk1Δ*, but because Ctk1-IS inhibition alone produced strong loss of Ser2P in our system, no additional effect of Bur1 inactivation could be seen (**Fig 4**). These results confirm our earlier conclusions that Ctk1/Cdk12 produces the vast majority of Ser2P (24, 35), although Bur1/Cdk9 may contribute a small amount of Ser2P early in elongation ((8), **Fig 3B**). Mutations in genes for Bur1 or its associated cyclin Bur2 exhibit phenotypes and genetic interactions diagnostic of elongation defects, so the partial drop in Ser2P may be explained, at least in part, by inefficient elongation leading to reduced levels of elongation complexes.

Our finding that Bur1/Cdk9 phosphorylates Rpb1 linker sites mediating binding of Spt6 further links this kinase to promotion of transcription elongation. The other well-characterized Bur1 substrate is the C-terminal repeat region of Spt5, where phosphorylation promotes binding of the PAF complex elongation factor (27, 36). Although some groups have reported that RNApII crosslinking to transcribed genes is not affected by deletion of *BUR2* (8, 37), it is hard to imagine how RNApII elongation could be normal given the very slow growth, elongation-related phenotypes, and defects in elongation factor recruitment in these cells. In contrast to these other reports, we observed a pronounced drop off in RNApII ChIP signal from 5’ to 3’ ends of genes in *bur2Δ* budding yeast (24). Similarly, recent experiments in S. pombe (17) showed that inhibition of a Cdk9-AS strain also caused a marked polar elongation defect. Finally, Cdk9 inhibition in mammalian cells blocks RNApII from escaping past the early pause site (38). Therefore, the preponderance of evidence indicates that Bur1/Cdk9 phosphorylates multiple substrates to promote efficient elongation.

In conclusion, we believe that covalent inhibition of the transcription-related CDKs can provide important information about their function. With the proper caveats and controls, the IS kinase alleles should prove to be useful tools for our future in vivo and in vitro experiments, and we look forward to distributing them to other labs for their work.

## Acknowledgements

We are grateful to Grant Hartzog (UC Santa Cruz) for anti-Spt5 serum, Dirk Eick (Munich) for phospho-CTD antibodies, Nathanael Gray and Nick Kwiatkowski for helpful discussions. This work was supported by NIH grants GM056663 to S.B., GM082545 to CPH, and GM116560 to CPH and TF.

## Methods

### Molecular genetics

Plasmids encoding HA3-tagged kinases were previously described (24, 39). Mutations were made using inverse PCR-mediated mutagenesis (primer sequences available upon request), and confirmed by DNA sequencing. Strains used in **Figures 1 and S1** were constructed using standard plasmid shuffling (40). For other figures, strains with kinase IS alleles integrated at the natural chromosomal locus were constructed using the *delitto perfetto* method (41). Correct clones were identified using the CMK sensitivity phenotype, followed by sequencing of PCR fragments amplified from the chromosomal locus. Yeast strains are listed in **Table 1**.

**Table 1.**
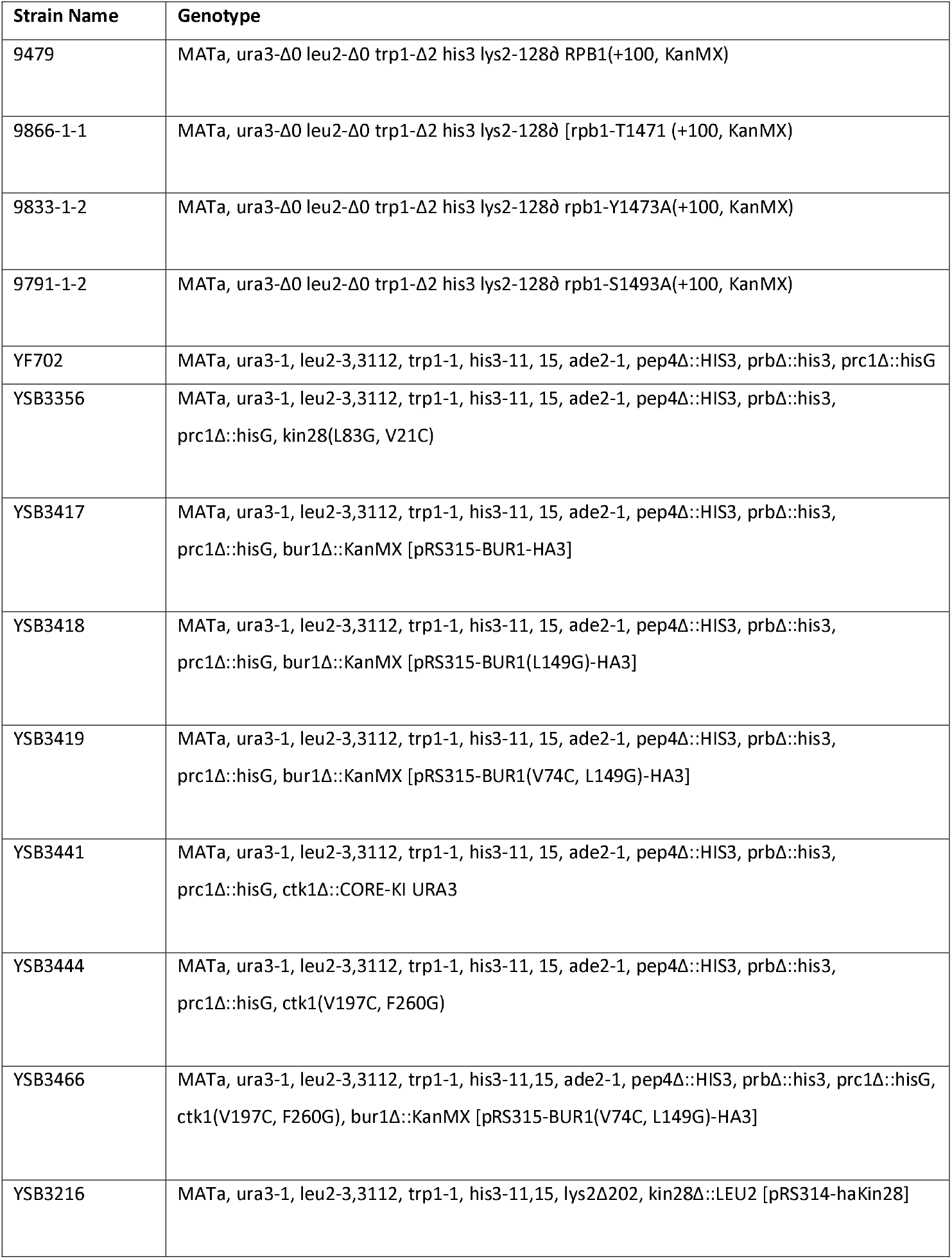
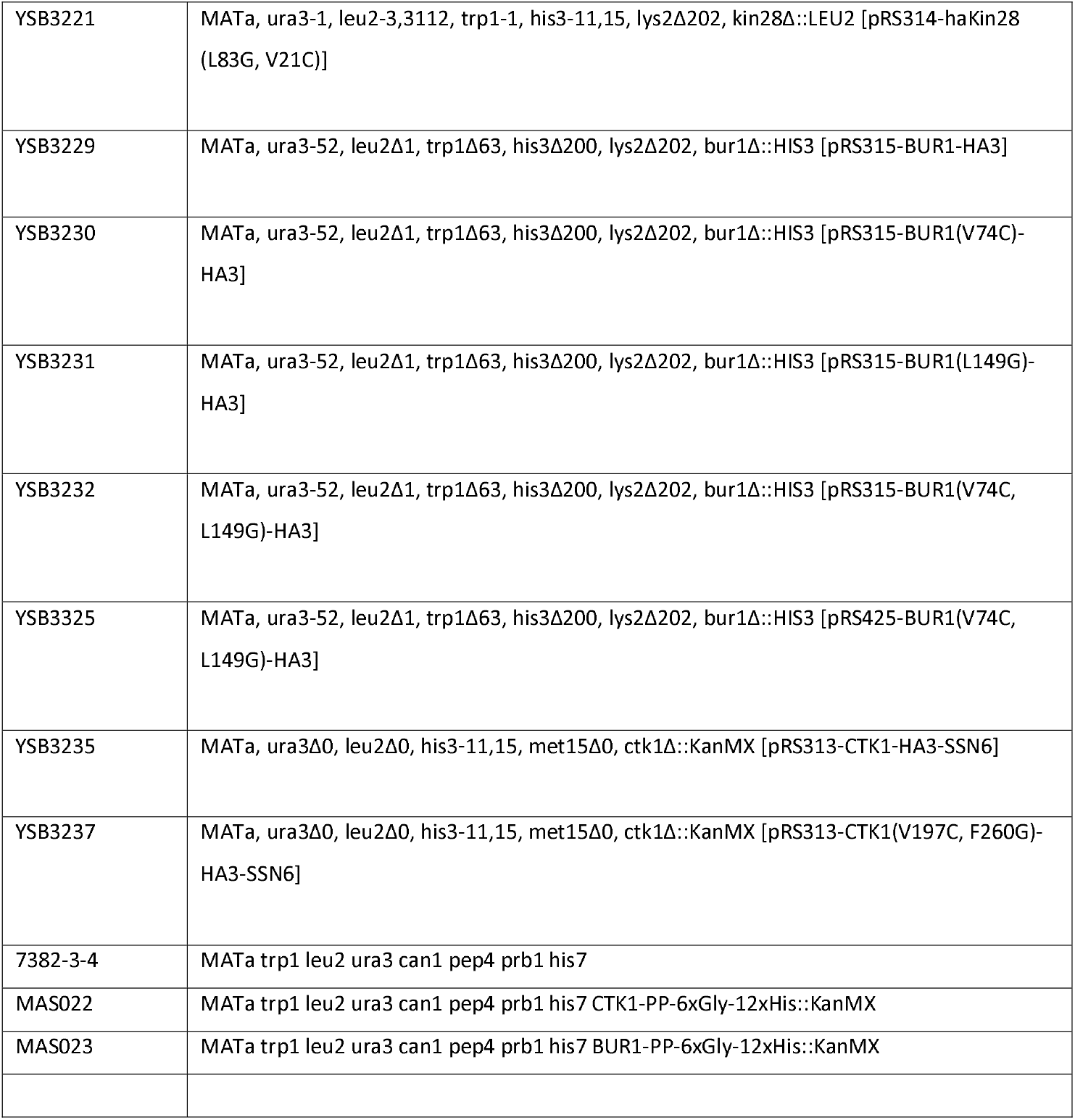
Yeast Strains used in this study.

### Immunoblotting

The indicated yeast strains were grown to logarithmic phase (OD595 of ~1.0) in standard YPD medium (except for **Figure 1**, which used synthetic complete media with appropriate omissions of amino acids for plasmid marker selection). Cells were collected by centrifugation and lysed with glass beads using either the TCA method (42) or lysis buffer [50 mM Tris-HCl (pH 8.0), 150 mM sodium chloride, 0.1% Nonidet-P40, supplemented with 1 mM PMSF, 1 μg/ml leupeptin, 1 μg/ml pepstatin A, and 1 μg/ml aprotinin) as described in (24, 39). Protein concentrations in lysates were determined using the BioRad DC or Pierce Coomassie Protein Assay reagents, and 30-50 μg of total proteins were separated by SDS-PAGE on 8% or 10% polyacrylamide gels, depending on the size of the proteins to be imaged. Proteins were transferred to nitrocellulose membranes and probed with the antibodies indicated. For **Sup Fig 2**, Rpb1 linker phosphorylations were assayed with anti-pT1471 (1:10,000), anti-pY1473 (1:500), anti-pS1493 (1:50,000), anti-Rpb1-CTD (8WG16, 1:5000) as described in (25). The secondary antibody was goat anti-rabbit serum coupled to an infrared tag (IR 800CW) for detection using a LiCor Odyssey. For other figures, monoclonal anti-phospho CTD antibodies were provided by Dirk Eick (6) (3E8, 3E10, 4D12, 4E12, 6G7; all used at 1:1000), generated in-house (H5, H14, 8WG16, TBP; all 1:1000, except 1:2500 for TBP), or purchased commercially (Rpb3 1Y26 from Neoclone, anti-HA 12013819001 from Roche, 1:1000). The secondary antibody was goat anti-rabbit (Sigma A0545) or goat anti-mouse (Jackson ImmunoResearch 115-035-044) antibodies coupled to horseradish peroxidase for detection on film using Pierce Supersignal West Chemiluminescent Substrate (Pico, except where noted when Femto Maximum Sensitivity was used).

### Antibodies

Polyclonal antiserum that specifically recognizes Rpb1 pS1493 was previously described (25). Polyclonal antiserum against Rpb1 pT1471 and pY1473 were produced by Covance. Briefly, rabbits UT765 and UT768 were injected with the Rpb1 linker-derived peptide CGQDGGVTP(pY)SNESGLVN conjugated to KLH and rabbits UT763 and UT764 were injected with the Rpb1 linker-derived peptide CEDGQDGGV(pT)PYSNESGL conjugated to KLH. The exsanguination bleed was subjected to positive and negative affinity purification steps over columns consisting of the phosphorylated peptide or unphosphorylated peptide, respectively. The specificity for the phosphorylated peptide versus the unphosphorylated peptide was validated by western blots using WT strains or mutants with alanine substitutions at the modification sites (**Sup Fig 2**).

### Peptide binding southwestern assay

His-MBP-Rpb1 (1456-1511) protein was diluted to 0.5 mg/mL in kinase buffer (25 mM HEPES pH 7.5, 50 mM potassium acetate, 10 mM MgCl2, 10% glycerol, 2 mM DTT, 0.5 mM ATP) and 80 μl aliquots were prepared. Kinase complexes were prepared as described below, and 1.5 μl of each kinase was added to the MPB-Rpb1 and incubated at 30°C for one hour. For blotting, 15 μL of reaction was transferred to a tube containing 5 μl 4x SDS-loading dye. 5 μL of each sample was loaded on a 12% gel and electrophoresed for 1 hour at 160 V. Far western blots were performed as described previously (25).

### Protein expression and purification

Wild type Ctk1 and Bur1 complexes were purified from Saccharomyces cerevisiae strains MAS022 and MAS023 respectively, containing a PreScission Protease cleavable His-12 tag fused to the target proteins. Eight liters of culture of each strain and a parallel culture of the parental strain 7382-3-4 lacking tagged kinases were grown in YPD (yeast extract peptone dextrose) inoculated with 50 mL of a saturated overnight culture and incubated at 30°C until they reached an OD of about 3. Cells were harvested by centrifugation, washed once with cold water, and pelleted by centrifugation. Cells were frozen by passing through a syringe into liquid nitrogen and lysed under liquid nitrogen using a SPEX SamplePrep 6870 Freezer/Mill (SPEX SamplePrep, Metuchen, NJ). Pulverized yeast were thawed in 2 pellet equivalents of lysis buffer (50 mM Tris-Cl pH 7.5, 1 M NaCl, 10% glycerol, 40 mM imidazole, 1.4 mg/mL pepstatin, 1 mg/mL leupeptin, 1 mg/mL aprotinin, 1.9 mM PMSF). Lysates were clarified by centrifugation at 37,000 RCF for 30 min. The supernatant was incubated with 0.5 mL of Ni-NTA agarose (Qiagen) for 30 min 4°C with agitation, followed by 3 washes with 5 mL of lysis buffer, then 2 washes with 5 mL of wash buffer (25 mM Tris-Cl pH 7.5, 150 mM NaCl, 10% glycerol, 40 mM imidazole) and finally eluted with 4 times 2 mL of elution buffer (25 mM Tris-Cl pH 7.5, 150 mM NaCl, 10% glycerol, 300 mM imidazole). The eluted protein was incubated with 20 mg PreScission Protease in 6,000-8,000 FisherbrandTM regenerated cellulose dialysis tubing in dialysis buffer (50 mM Tris-Cl pH 7.5, 500 mM NaCl, 10% glycerol, 15 mM imidazole, 2 mM BME) overnight. The protease was removed with glutathione-agarose. The subsequent flowthrough fraction was collected, and the resin was washed with 2 times 1 mL of dialysis buffer. The flow through and wash were pooled and concentrated to 200 μL using a 30 kDa Vivaspin concentrator (Sartorius Stedim Biotech, Aubagne, France) and flash frozen in liquid nitrogen prior to storage at 80°C. The negative control strain was processed in parallel. Coomassie stained gels of each preparation is shown in **Sup Fig 2B**.

GST-Spt6 tSH2 (residues 1247-1451) was expressed and purified as described previously (25), and are shown in **Sup Fig 2A**. His-MBP-Rpb1 (residues 1456-1511) and mutated derivatives were expressed from pET17B vectors (Invitrogen) in BL21 codon plus (RIL) E. coli cells (Stratagene). Cultures were grown in 2 L of autoinduction media (43)in baffled 1.8 L flasks at 37°C with continuous shaking. After 8 hours, the cultures were shifted to 19°C and shaken for an additional 16–24 hours. Harvested cells were stored at −80°C. Cells were thawed and lysed in buffer containing lysozyme, DNase and protease inhibitors, followed by sonication. Lysates were clarified by centrifugation at 15,000 rpm for 35 min. The supernatant was incubated with 6 mL of Ni-NTA agarose (Qiagen) for 30 min at 4°C with agitation, followed by 3 washes with 25 mL of lysis buffer, then 2 washes with 15 mL of wash and finally eluted with 2 times 20 mL of elution buffer.

## Supplemental Figure Legends

**Sup Fig 1.**
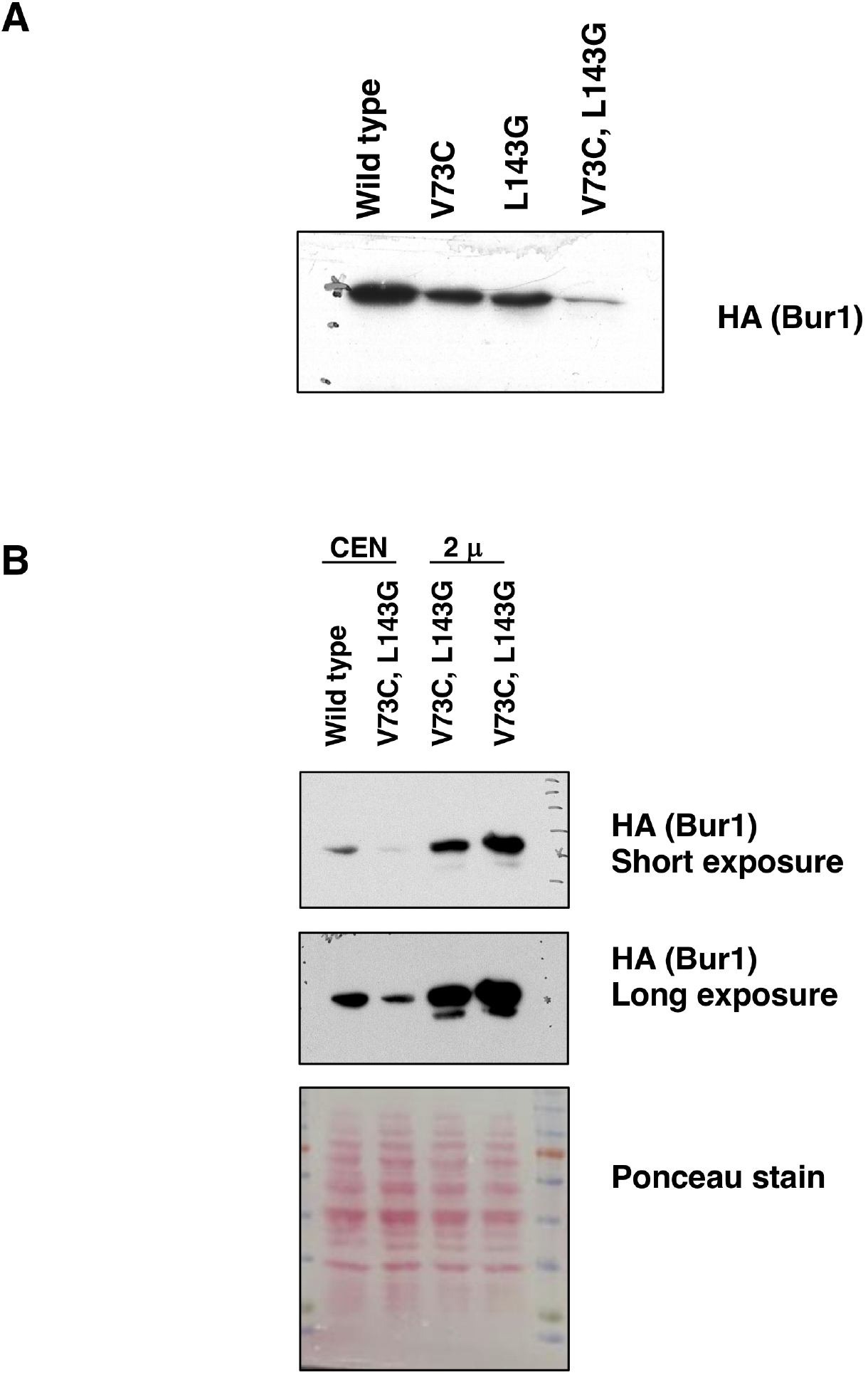
Bur1 IS mutations affect protein levels. A. Strains carrying the indicated Bur1 single or double mutations were assayed for protein levels by probing for the HA-epitope tag. Strains used are YSB3229 (WT), YSB3230 (V74C), YSB3231 (L149G), and YSB3232 (V74C, L149G). B. Levels of Bur1-IS protein can be boosted by high copy expression. Cell extracts were analyzed as in part A. Strains used are YSB3229 (WT), YSB3232 (V74C, L149G, CEN), and two isolates of YSB3325 (V74C, L149G, 2μ). In addition to two different exposures, a photograph of the Ponceau-stained blot is shown as a loading control.

**Sup Fig 2.**
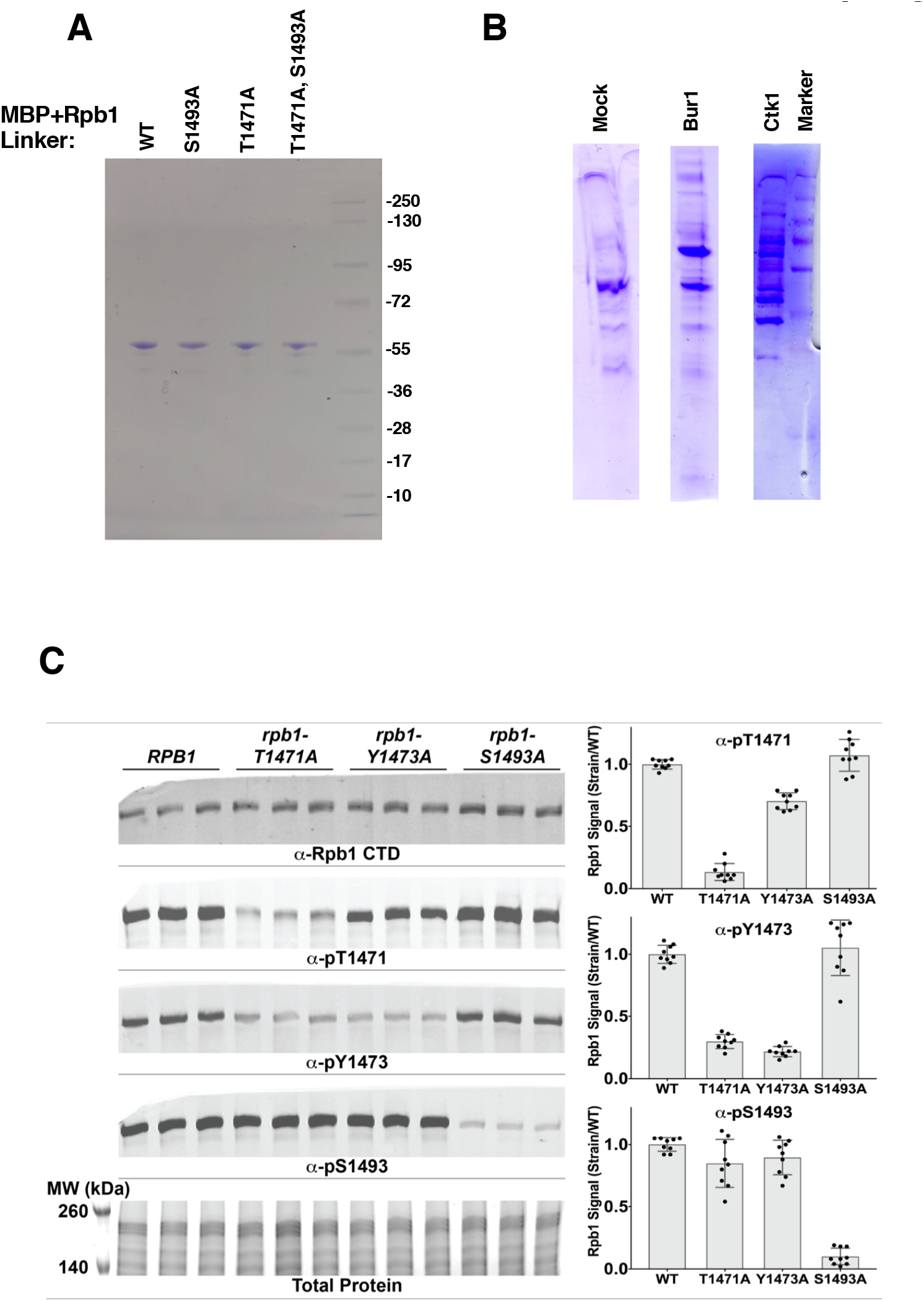
Specificity of antibodies against Rpb1 linker region phosphorylations. Three independent cultures of WT (Rpb1) or mutant yeast strains with alanine substitutions at the three mapped Rpb1 linker phosphorylation sites (T1471, Y1473, and S1493) were tested with each phosphorylation-specific antibody. An antibody against the Rpb1 CTD and total protein staining are shown as loading controls. Signals from multiple experiments were quantified, normalized to the average signal from WT controls run on the same gel, then compiled in the panels on the right with the average and standard deviation shown. The rpb1-T1471A strain displayed reduced signal for phosphorylated T1471 as expected, but also produced less signal with the antibodies against phosphorylated Y1473, suggesting this threonine may be important for phosphorylation of Y1473. A similar but smaller effect was also seen with reduced phosphoT1471 signal in an rpb1-Y1473A strain. These changes could also reflect changes in the epitopes recognized by the antibodies.

## REFERENCES

1. Bishop AC, Ubersax JA, Petsch DT, Matheos DP, Gray NS, Blethrow J, Shimizu E, Tsien JZ, Schultz PG, Rose MD, Wood JL, Morgan DO, Shokat KM. 2000. A chemical switch for inhibitor-sensitive alleles of any protein kinase. Nature 407:395–401.

2. Cohen MS, Zhang C, Shokat KM, Taunton J. 2005. Structural bioinformatics-based design of selective, irreversible kinase inhibitors. Science 308:1318–1321.

3. Snead JL, Sullivan M, Lowery DM, Cohen MS, Zhang C, Randle DH, Taunton J, Yaffe MB, Morgan DO, Shokat KM. 2007. A coupled chemical-genetic and bioinformatic approach to Pololike kinase pathway exploration. Chem Biol 14:1261–1272.

4. Rodríguez-Molina JB, Tseng SC, Simonett SP, Taunton J, Ansari AZ. 2016. Engineered Covalent Inactivation of TFIIH-Kinase Reveals an Elongation Checkpoint and Results in Widespread mRNA Stabilization. Mol Cell 63:433–444.

5. Buratowski S. 2003. The CTD code. Nat Struct Biol 10:679–680.

6. Eick D, Geyer M. 2013. The RNA polymerase II carboxy-terminal domain (CTD) code. Chem Rev 113:8456–8490.

7. Corden JL. 2013. RNA polymerase II C-terminal domain: Tethering transcription to transcript and template. Chem Rev 113:8423–8455.

8. Qiu H, Hu C, Hinnebusch AG. 2009. Phosphorylation of the Pol II CTD by KIN28 enhances BUR1/BUR2 recruitment and Ser2 CTD phosphorylation near promoters. Mol Cell 33:752–762.

9. Liu Y, Kung C, Fishburn J, Ansari AZ, Shokat KM, Hahn S. 2004. Two cyclin-dependent kinases promote RNA polymerase II transcription and formation of the scaffold complex. Mol Cell Biol 24:1721–1735.

10. Kanin EI, Kipp RT, Kung C, Slattery M, Viale A, Hahn S, Shokat KM, Ansari AZ. 2007. Chemical inhibition of the TFIIH-associated kinase Cdk7/Kin28 does not impair global mRNA synthesis. Proceedings of the National Academy of Sciences 104:5812–5817.

11. Viladevall L, St Amour CV, Rosebrock A, Schneider S, Zhang C, Allen JJ, Shokat KM, Schwer B, Leatherwood JK, Fisher RP. 2009. TFIIH and P-TEFb coordinate transcription with capping enzyme recruitment at specific genes in fission yeast. Mol Cell 33:738–751.

12. Tietjen JR, Zhang DW, Rodríguez-Molina JB, White BE, Akhtar MS, Heidemann M, Li X, Chapman RD, Shokat K, Keles S, Eick D, Ansari AZ. 2010. Chemical-genomic dissection of the CTD code. Nat Struct Mol Biol 17:1154–1161.

13. Qiu H, Hu C, Gaur NA, Hinnebusch AG. 2012. Pol II CTD kinases Bur1 and Kin28 promote Spt5 CTR-independent recruitment of Paf1 complex. EMBO J 31:3494–3505.

14. Bataille AR, Jeronimo C, Jacques P-E, Laramée L, Fortin M-È, Forest A, Bergeron M, Hanes SD, Robert F. 2012. A universal RNA polymerase II CTD cycle is orchestrated by complex interplays between kinase, phosphatase, and isomerase enzymes along genes. Mol Cell 45:158–170.

15. Mbogning J, Pagé V, Burston J, Schwenger E, Fisher RP, Schwer B, Shuman S, Tanny JC. 2015. Functional interaction of Rpb1 and Spt5 C-terminal domains in co-transcriptional histone modification. Nucleic Acids Res 43:9766–9775.

16. Parua PK, Booth GT, Sansó M, Benjamin B, Tanny JC, Lis JT, Fisher RP. 2018. A Cdk9-PP1 switch regulates the elongation-termination transition of RNA polymerase II. Nature 558:460–464.

17. Booth GT, Parua PK, Sansó M, Fisher RP, Lis JT. 2018. Cdk9 regulates a promoter-proximal checkpoint to modulate RNA polymerase II elongation rate in fission yeast. Nat Commun 9:679.

18. Glover-Cutter K, Larochelle S, Erickson B, Zhang C, Shokat K, Fisher RP, Bentley DL. 2009. TFIIH-associated Cdk7 kinase functions in phosphorylation of C-terminal domain Ser7 residues, promoter-proximal pausing, and termination by RNA polymerase II. Mol Cell Biol 29:5455–5464.

19. Larochelle S, Amat R, Glover-Cutter K, Sansó M, Zhang C, Allen JJ, Shokat KM, Bentley DL, Fisher RP. 2012. Cyclin-dependent kinase control of the initiation-to-elongation switch of RNA polymerase II. Nat Struct Mol Biol 19:1108–1115.

20. Ebmeier CC, Erickson B, Allen BL, Allen MA, Kim H, Fong N, Jacobsen JR, Liang K, Shilatifard A, Dowell RD, Old WM, Bentley DL, Taatjes DJ. 2017. Human TFIIH Kinase CDK7 Regulates Transcription-Associated Chromatin Modifications. Cell Rep 20:1173–1186.

21. Kwiatkowski N, Zhang T, Rahl PB, Abraham BJ, Reddy J, Ficarro SB, Dastur A, Amzallag A, Ramaswamy S, Tesar B, Jenkins CE, Hannett NM, McMillin D, Sanda T, Sim T, Kim ND, Look T, Mitsiades CS, Weng AP, Brown JR, Benes CH, Marto JA, Young RA, Gray NS. 2014. Targeting transcription regulation in cancer with a covalent CDK7 inhibitor. Nature 511:616–620.

22. Olson CM, Jiang B, Erb MA, Liang Y, Doctor ZM, Zhang Z, Zhang T, Kwiatkowski N, Boukhali M, Green JL, Haas W, Nomanbhoy T, Fischer ES, Young RA, Bradner JE, Winter GE, Gray NS. 2018. Pharmacological perturbation of CDK9 using selective CDK9 inhibition or degradation. Nat Chem Biol 14:163–170.

23. Laitem C, Zaborowska J, Isa NF, Kufs J, Dienstbier M, Murphy S. 2015. CDK9 inhibitors define elongation checkpoints at both ends of RNA polymerase II-transcribed genes. Nat Struct Mol Biol 22:396–403.

24. Keogh MC, Podolny V, Buratowski S. 2003. Bur1 kinase is required for efficient transcription elongation by RNA polymerase II. Mol Cell Biol 23:7005–7018.

25. Sdano MA, Fulcher JM, Palani S, Chandrasekharan MB, Parnell TJ, Whitby FG, Formosa T, Hill CP. 2017. A novel SH2 recognition mechanism recruits Spt6 to the doubly phosphorylated RNA polymerase II linker at sites of transcription. Elife 6:213.

26. Vos SM, Farnung L, Boehning M, Wigge C, Linden A, Urlaub H, Cramer P. 2018. Structure of activated transcription complex Pol II-DSIF-PAF-SPT6. Nature 560:607–612.

27. Liu Y, Warfield L, Zhang C, Luo J, Allen J, Lang WH, Ranish J, Shokat KM, Hahn S. 2009. Phosphorylation of the transcription elongation factor Spt5 by yeast Bur1 kinase stimulates recruitment of the PAF complex. Mol Cell Biol 29:4852–4863.

28. Joo YJ, Ficarro SB, Chun Y, Marto JA, Buratowski S. In vitro analysis of RNA polymerase II elongation complex dynamics. Bioarxiv.

29. Suh H, Ficarro SB, Kang U-B, Chun Y, Marto JA, Buratowski S. 2016. Direct Analysis of Phosphorylation Sites on the Rpb1 C-Terminal Domain of RNA Polymerase II. Mol Cell 61:297–304.

30. Nilson KA, Guo J, Turek ME, Brogie JE, Delaney E, Luse DS, Price DH. 2015. THZ1 Reveals Roles for Cdk7 in Co-transcriptional Capping and Pausing. Mol Cell 59:576–587.

31. Zeng M, Kwiatkowski NP, Zhang T, Nabet B, Xu M, Liang Y, Quan C, Wang J, Hao M, Palakurthi S, Zhou S, Zeng Q, Kirschmeier PT, Meghani K, Leggett AL, Qi J, Shapiro GI, Liu JF, Matulonis UA, Lin CY, Konstantinopoulos PA, Gray NS. 2018. Targeting MYC dependency in ovarian cancer through inhibition of CDK7 and CDK12/13. Elife 7: e39030.

32. Søgaard TMM, Svejstrup JQ. 2007. Hyperphosphorylation of the C-terminal repeat domain of RNA polymerase II facilitates dissociation of its complex with mediator. Journal of Biological Chemistry 282:14113–14120.

33. Jones JC, Phatnani HP, Haystead TA, MacDonald JA, Alam SM, Greenleaf AL. 2004. C-terminal repeat domain kinase I phosphorylates Ser2 and Ser5 of RNA polymerase II C-terminal domain repeats. Journal of Biological Chemistry 279:24957–24964.

34. Bartkowiak B, Greenleaf AL. 2014. Expression, Purification, and Identification of Associated Proteins of the Full Length hCDK12/CyclinK Complex. Journal of Biological Chemistry 290: 1785-1795.

35. Cho EJ, Kobor MS, Kim M, Greenblatt J, Buratowski S. 2001. Opposing effects of Ctk1 kinase and Fcp1 phosphatase at Ser 2 of the RNA polymerase II C-terminal domain. Genes Dev 15:3319–3329.

36. Zhou K, Kuo WHW, Fillingham J, Greenblatt JF. 2009. Control of transcriptional elongation and cotranscriptional histone modification by the yeast BUR kinase substrate Spt5. Proc Natl Acad Sci USA 106:6956–6961.

37. Chu Y, Simic R, Warner MH, Arndt KM, Prelich G. 2007. Regulation of histone modification and cryptic transcription by the Bur1 and Paf1 complexes. EMBO J 26:4646–4656.

38. Jonkers I, Kwak H, Lis JT. 2014. Genome-wide dynamics of Pol II elongation and its interplay with promoter proximal pausing, chromatin, and exons. 3:e02407.

39. Keogh MC, Cho E-J, Podolny V, Buratowski S. 2002. Kin28 is found within TFIIH and a Kin28-Ccl1-Tfb3 trimer complex with differential sensitivities to T-loop phosphorylation. Mol Cell Biol 22:1288–1297.

40. Guthrie C, Fink GR. 1991. Guide to yeast genetics and molecular biology. Methods in Enzymology. vol. 194. Academic Press, New York, NY.

41. Storici F, Resnick MA. 2006. The delitto perfetto approach to in vivo site-directed mutagenesis and chromosome rearrangements with synthetic oligonucleotides in yeast. Meth Enzymol 409:329–345.

42. McCullough L, Connell Z, Petersen C, Formosa T. 2015. The Abundant Histone Chaperones Spt6 and FACT Collaborate to Assemble, Inspect, and Maintain Chromatin Structure in Saccharomyces cerevisiae. Genetics 201:1031–1045.

43. Studier FW. 2005. Protein production by auto-induction in high density shaking cultures. Protein Expr Purif 41:207–234.

